# Quantifiable TCR repertoire changes in pre-diagnostic blood specimens among high-grade ovarian cancer patients

**DOI:** 10.1101/2023.10.12.562056

**Authors:** Xuexin Yu, Jianfeng Ye, Cassandra A. Hathaway, Shelley Tworoger, Jayanthi Lea, Bo Li

**Author notes:** Corresponding authors: Bo Li and Jayanthi Lea.

## Abstract

High grade serous ovarian cancer (HGOC) is a major cause of death in women. Early detection of HGOC usually leads to a cure, yet it remains a clinical challenge with over 90% HGOCs diagnosed at advanced stages. This is mainly because conventional biomarkers are not sensitive to detect the microscopic yet metastatic early HGOC lesions. In this study, we sequenced the blood T cell receptor (TCR) repertoires of 466 ovarian cancer patients and controls, and systematically investigated the immune repertoire signatures in HGOCs. We observed quantifiable changes of selected TCRs in HGOCs that are reproducible in multiple independent cohorts. Importantly, these changes are stronger during stage I. Using pre-diagnostic patient blood samples from the Nurses’ Health Study, we confirmed that HGOC signals can be detected in the blood TCR repertoire up to 4 years proceeding conventional diagnosis. Our findings may provide the basis of an immune-based HGOC early detection criterion.

**Statement of significance:** We made an unprecedented discovery that a strong and quantifiable change in the blood TCR repertoire occurs 4 to 2 years before high grade ovarian cancers could be diagnosed with conventional clinical tests. This finding might be useful to develop novel screening biomarkers to detect early-stage ovarian cancers.

## Introduction

Ovarian cancer accounts for 2.5% of all malignancies seen in women^1^, while causing 5% of cancer-related deaths^2^. High-grade serous ovarian carcinoma (HGOC), which comprise over 70% of incident ovarian tumors^1^, contributes to the high mortality burden of this disease, whereas low-grade ovarian tumors are much less lethal^3^. Stage I high grade serous carcinomas are confined within ovaries or fallopian tubes, and are largely curable with complete surgical resection and chemotherapy (93% 5-year relative survival)^4^. In contrast, HGOCs diagnosed at advanced stages have a 5-year survival of 31%^4^. HGOCs are believed to arise from a range of epithelial changes with p53 mutations, serous tubal intraepithelial carcinoma (STIC) in the fallopian tube and atypical lesions inbetween p53 mutations and STIC^5^. The current paradigms of serous carcinogenesis include a precursor lesion (STIC) with gradual progression to cancer and precursor metastasis into the peritoneal cavity^6,7^. All paradigms allude detection or prevention by conventional methods and consequently, HGOCs are rarely found at stage I, with over 87% cases diagnosed at stage III or IV^1^.

Given the apparent clinical benefit of detecting ovarian cancer early, noninvasive assays have been evaluated in large-scale, prospective screening trials, largely focusing on serum CA-125 levels^8^ and its changes over time^9^, serum human epididymis protein 4 (HE-4)^10,11^ and transvaginal ultrasound^12^. However, a recent longitudinal trial of over 200,000 subjects followed for more than 18 years showed no mortality benefit for HGOC among women routinely tested by one or a combination of these assays^13^. Studies have demonstrated that most conventional biomarkers have limited predictive ability until 6-12 months before diagnosis^14^, possibly because HGOC primarily comprises of microscopic lesions until very late in its progression. As such, blood biomarkers or tumor imaging may not be sensitive to detect the early-stage ovarian tumors.

Previously, we showed that early-stage cancers induce observable changes in the blood T cell receptor (TCR) repertoire, providing an alternative for noninvasive cancer diagnosis^15^. Although it remains unclear what causes these conservative changes in the TCR repertoires of patients with diverse genetic backgrounds, the concept of immunoediting^16,17^ may explain why signals can be seen at early stages. Specifically, during the ‘Elimination’ phase, exposure to early tumor antigens could result in a rapid expansion of cancer-associated T cells^18^, leading to detectable signals in the TCR repertoire in circulating white blood cells. Here, we first developed a new method to quantitatively dissect the TCR repertoire data into quantifiable functional units. We then collected preoperative blood samples from patients with ovarian tumors to identify TCR biomarkers that are enriched in the HGOC patients compared to women with benign ovarian tumors. Finally, we measured TCRs in pre-diagnostic blood specimens from patients diagnosed with ovarian cancer within 5 year after blood draw using samples from a large longitudinal cohort study and matched controls. Our analysis revealed transient but significant TCR repertoire changes that occurred up to 4 years prior to conventional ovarian cancer diagnosis.

## Results

### Trimer embedding of T cell receptors and RFU definition

We first obtained a numeric embedding of the CDR3β region that preserved sequence similarity (Figure 1a). In brief, approximately 20 million TCRs from the public domain (Table S1) were clustered based on the variable gene (TRBV) and CDR3β sequences (Figure 1b) by GIANA^19^ to construct a trimer substitution matrix (Figure 1c-d). Approximated isometric embedding of each trimer was obtained using multidimensional scaling (Figure 1e) and the final embedding vector for each CDR3β was calculated by mean pooling of all consecutive trimers in the amino acid sequence. We benchmarked this embedding using 1,031 TCRs with known specificity to 10 common immunogenic epitopes^20^ (Table S2). Specifically, we obtained the numeric embedding of each TCR and calculated the Euclidean distances for each pair of TCRs. This distance was used to predict if the pair of TCRs were specific to the same antigen, and reached an Area under the Receiver Operative Characteristic curve (AUC) of 0.64 (Figure S1a-b). At a high specificity of 0.95, this method reached a sensitivity of 0.22, comparable to the state-of-art methods based on TCR similarity^19,21–23^. This result indicated that trimer embedding method preserves the ‘local specificity’ of TCRs, *i.e.* if the distance of two TCRs continuously decreases to 0, the probability that they share antigen-specificity will approach to 1. This property is guarded by the fact that TCR sequence similarity can be used as a surrogate for antigen specificity^21^.

**Figure 1.**
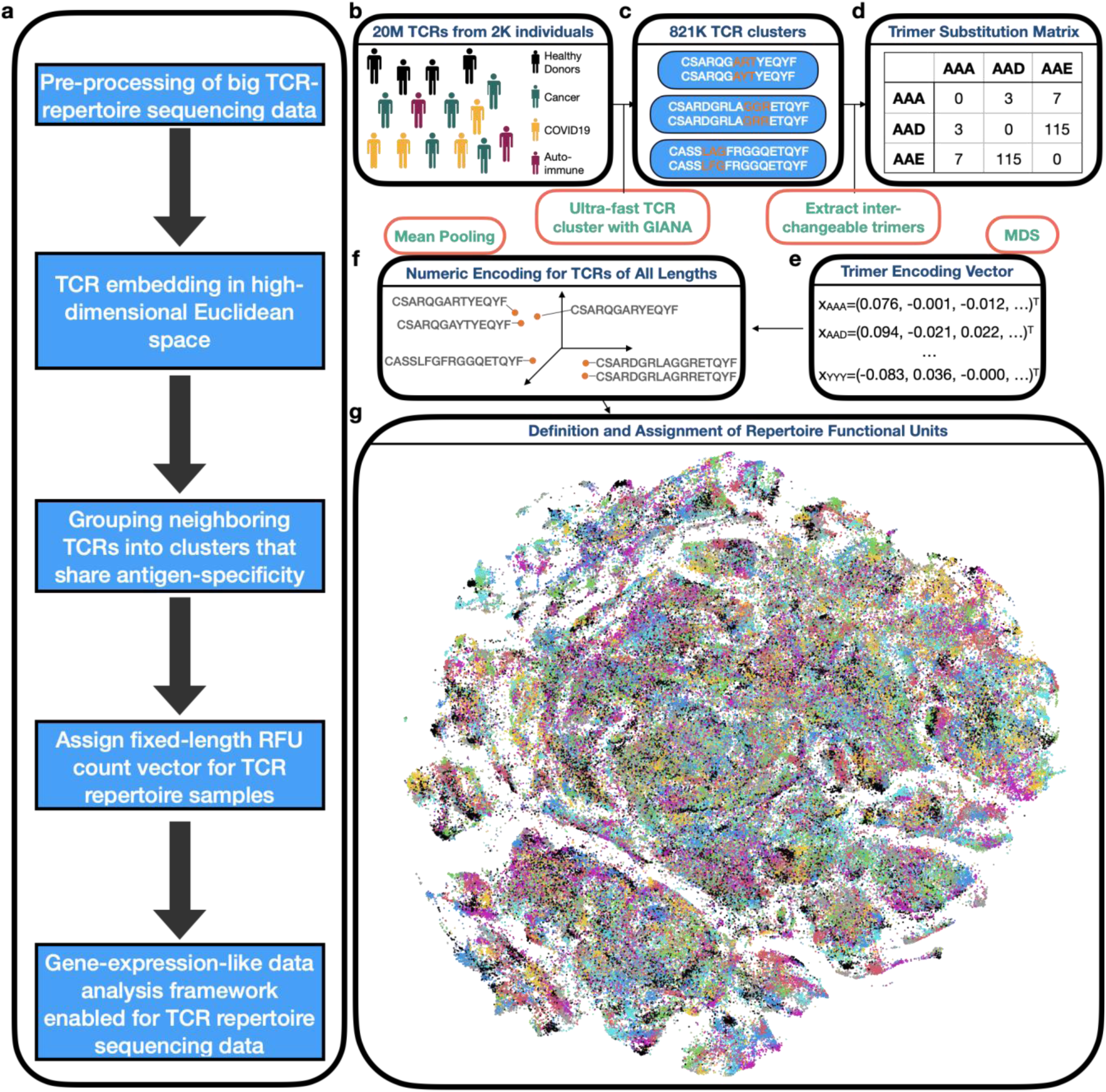
Trimer-guided embedding for TCRs and derivation of Repertoire Functional Units (RFU). a) Method workflow. The first 3 steps describe the trimer-embedding Euclidean space and last two steps describe how RFUs are defined. b) Massive clustering of TCRs from patients with diverse health conditions based on CDR3 amino acid sequence similarity. c) Illustration of replaceable trimers from small TCR clusters. d) Illustration of the trimer substitution matrix with each number represent the times a row trimer is replaced by the column trimer in a TCR cluster. e) Derivation of approximately isometric embedding for each trimer based on multidimensional scaling from the trimer substitution matrix in d). f) Representation of each CDR3 sequence in the high-dimensional Euclidean space by averaging all the consecutive trimers. g) RFU definition by pooling 1.2 million TCRs from 120 individuals shown as t-SNE plot. Colors denote distinct clusters with cluster centroids assigned by k-means.

With this property, we defined the ‘neighborhood’ in the TCR space, as local TCR clusters that likely recognize the same antigens. Such neighborhoods may carry disease-specific information. For example, a simple comparison between a B cell lymphoma sample and healthy control^24^ revealed several TCR neighborhoods enriched or depleted in the lymphoma patient, each characterized by conserved CDR3 motifs (Figure S1c-d). Systematic investigation by pooling over 1 million TCRs from 120 healthy donors^25^ revealed conservative TCR clusters seen in multiple individuals (Figure S1e). We thus divided the TCR space into 5,000 groups (Figure 1g), with over 84% of TCRs within 0.018 distance to the centroid, which is the cutoff of 90% specificity in the benchmark (Figure S1f). The group centroid was defined as a ‘Repertoire Functional Unit’, or RFU, which can be viewed as the ‘gene’ of a repertoire in the sense of antigen recognition. This definition allowed us to transform each TCR repertoire sample into a fixed-length numeric vector, *i.e.* the normalized TCR count of each RFU.

### TCR repertoire landscape in high-grade ovarian cancer patients

We prospectively collected a discovery cohort of preoperative PBMC samples from 213 women, including 67 patients with high-grade serous, 49 with other histologies and 97 with benign ovarian tumors. TCR repertoire sequencing data was obtained for each sample. Despite attempt to frequency match on age, cancer patients were significantly older than benign controls (Table S3). Therefore, we first investigated the impact of age over RFUs in the healthy individuals using publicly available TCR-seq cohorts. We first analyzed the Emerson et al. cohort^25^, which contained blood TCR repertoires of 666 healthy donors collected before 2017. Our analysis revealed that a subset of RFUs showed strong negative correlation with age (Figure S2a), which was further confirmed using another cohort^26^ (Figure S2b). The second cohort was comprised of 1,414 COVID-19+ individuals collected in 2020, which were also considered as healthy controls. Importantly, the RFUs with strong age associations in both cohorts were highly reproducible (Figure S2c). These results suggested that age-associated RFUs are conserved in the general population, providing the rationale to exclude age as a confounder in our downstream analysis.

We then visualized the TCR repertoires of all 213 individuals using the top 1,500 most variable RFUs (ranked by standard deviation). Unsupervised hierarchical clustering revealed a distinguishable separation between HGOCs and benign samples (Figure 2a), suggesting global difference in the immune repertoire between these two conditions. Principal component analysis (PCA) of the RFU matrix confirmed that PC1 is driven by disease categories (Figure 2b-c). In contrast, PC2 is mainly influenced by race, with African American patients showing the largest separation from Asian patients (Figure 2d-e). To systematically investigate the differences of TCR repertoire between HGOC and benign patients, we performed logistic regression adjusted for patient age and race for all 1,500 RFUs and observed significant results at FDR>0.2 (Figure 2f). In contrast, comparison between HGOC vs low-grade and low-grade vs benign yielded no significant RFUs, potentially due to limited sample size (Figure S2d). Next, we visualized the CDR3 motifs of the top up-/down-regulated ones in HGOC patients (Figure 2g). We noted a conservative “RLAG” pattern at the 6-9^th^ positions of RFU 1804. CDR3s with this pattern, combined with the use of joining gene TRBJ2-3*01 (DTQYF) have been reported to recognize the ELAGIGLTV epitope from melanoma antigen *MART-1*^27^, which is reportedly expressed in ovarian neoplasms^28^.

**Figure 2.**
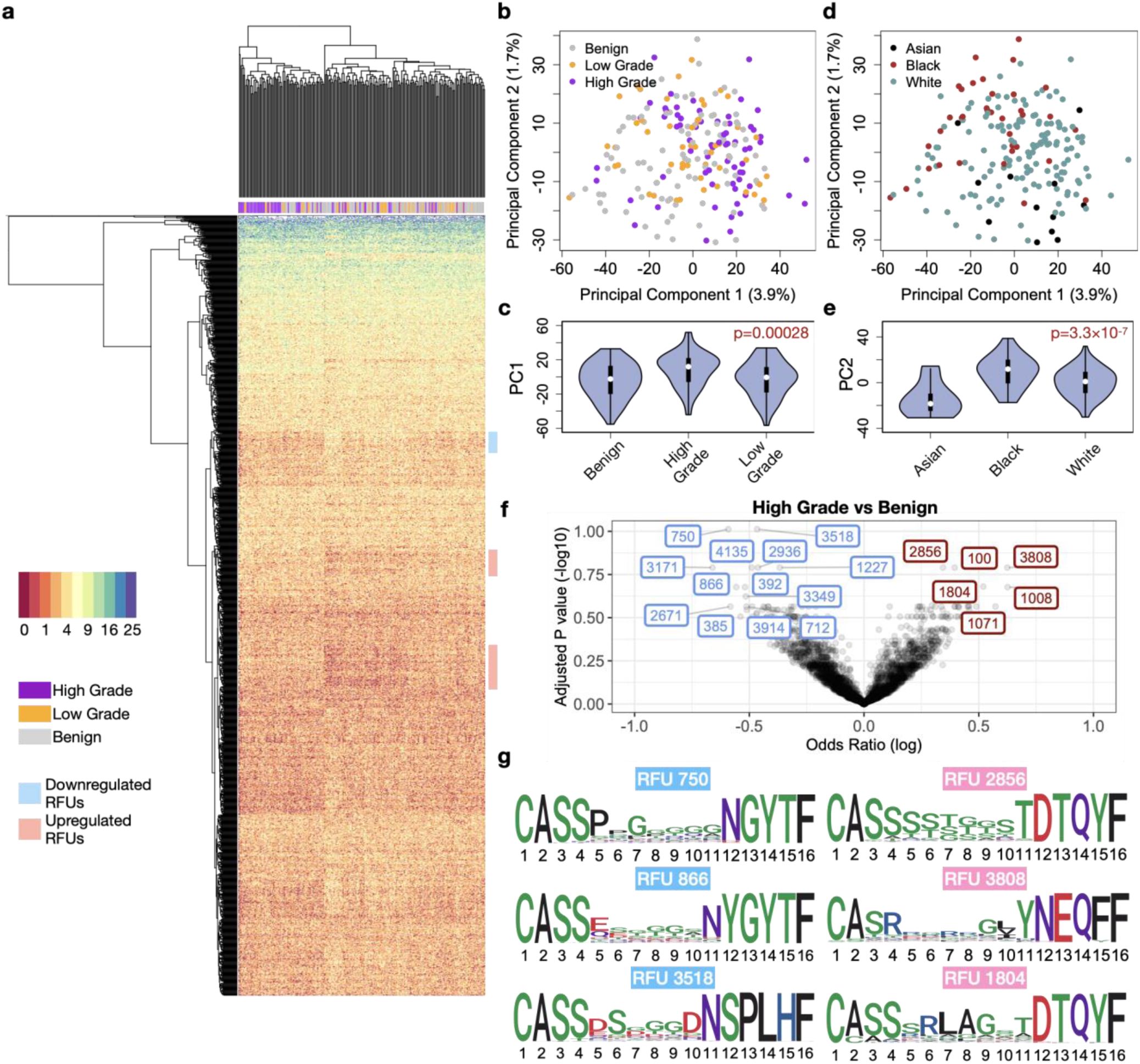
Characterization of TCR repertoire landscape in ovarian cancer patients. a) Heatmap showing the distribution of the top 1,500 most variable RFUs of high-grade, low-grade and benign patients. b) Distribution of ovarian cancer and benign patients on the PCA plot calculated from the RFU-by-patient matrix. c) Violin plot showing the differences of PC1 across disease categories. Statistical significance was evaluated using one-way ANOVA. d) Distribution of patient races on the PCA plot. e) Violin plot showing the differences of PC2 across patient races. Statistical significance was evaluated using one-way ANOVA. f) Volcano plot showing the log odds ratio vs FDR adjusted by Benjamini-Hochberg method. Odds ratio is estimated from logistic regression with disease status as a binary outcome, with each RFU being the covariate and adjusted for age and race. Blue: downregulated; Red: upregulated. g) Sequence logo analysis of selected top up-/down-regulated RFUs.

### RFU as a risk marker for HGOC

The above results indicated that selected RFUs are significantly altered in the blood repertoire of HGOCs compared to benign controls. We therefore proceeded to select a subset of RFUs to evaluate the risk of HGOC. First, we observed that although some RFUs reached high odds ratios in the logistic regression, there was no difference in the median levels of these RFUs between HGOC and benign patients (Figure 3a), suggesting the influence of outliers. After removal of such RFUs, we then defined the top 2 up- and down-regulated RFUs based on odds ratios, which included RFUs 750, 866, 3808 and 1804. Since none of these RFUs were age-related, we no longer considered age as a confounder in the following analysis (Figure S3a). We performed a survey of these RFUs across a wide spectrum of human cancers using blood repertoire samples (Table S1). Interestingly, in addition to HGOC, RFU 750 was also downregulated in melanoma and kidney cancer, where RFU 866 was only downregulated in HGOCs (Figure 3b). On the other hand, RFU 1804 is also upregulated in lung cancer, while RFU 3808 was higher in head and neck cancer (Figure S3b). Notably, for all four RFUs, healthy control samples from both children and adult cohorts had similar distributions as benign patients.

**Figure 3.**
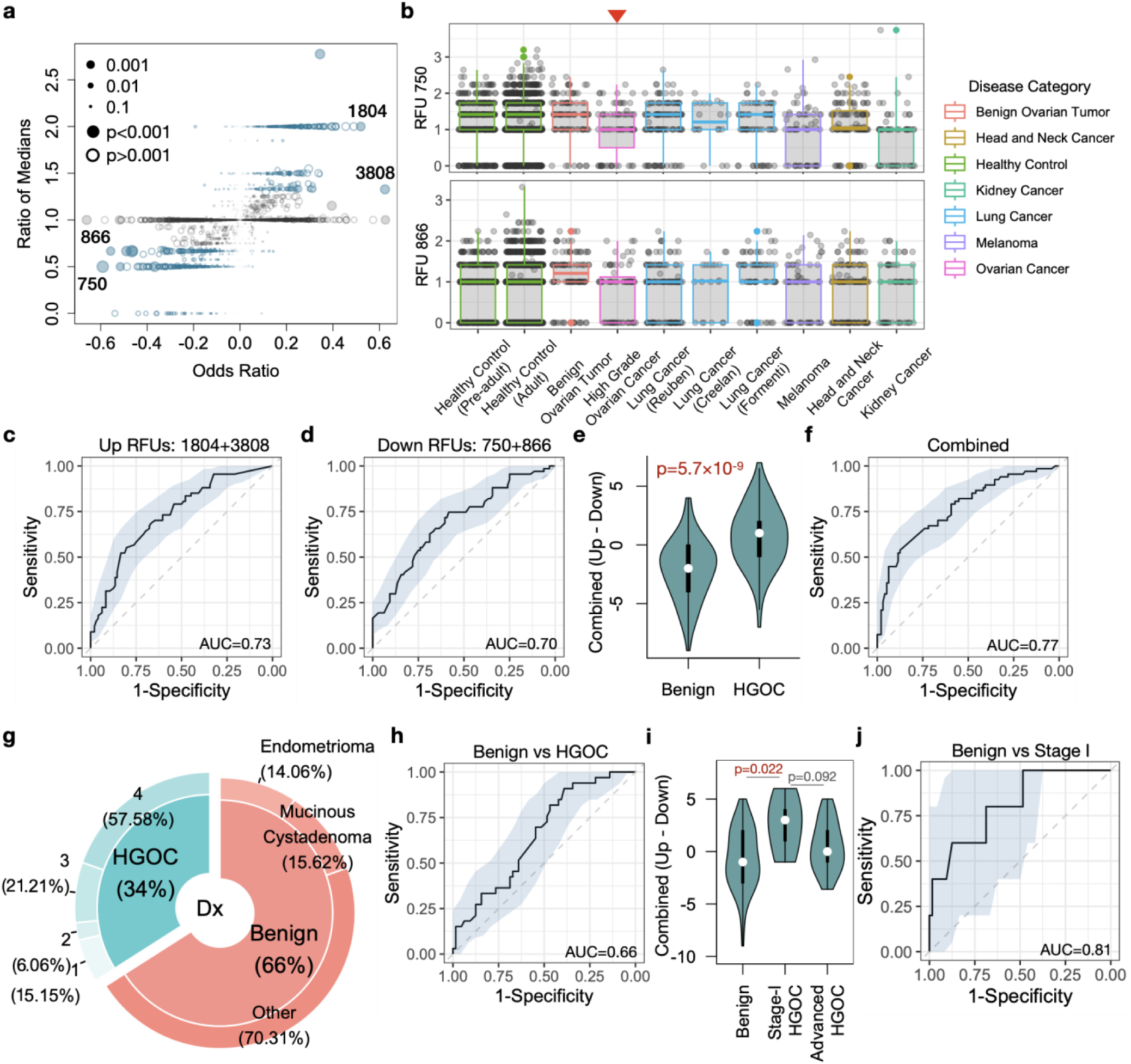
Selected RFUs as biomarker to distinguish HGOCs from benign ovarian lesions. a) Selection criteria for the top informative RFUs. Odds ratios and ratio of medians were described in Results. Blue color indicates ratio median <0.7 or >1.3. b) Boxplot showing the distributions of selected RFUs across multiple cancer types. All analysis was performed using blood TCR repertoire samples from the public domain. c-d) ROC curves showing the prediction accuracy of up- or down-regulated RFUs to predict HGOCs from benign patients. e) Combination of up- and down-regulated RFUs as a joint biomarker, OV RFU score. Statistical significance was evaluated using two-sided Wilcoxon test. g) Donut plot showing the sample composition in the validation cohort, with total N=97. h) Performance of OV RFU score in the validation cohort. i) Violin plot showing the distributions of OV RFU scores across benign, stage-I HGOC and advanced HGOCs. Statistical significance was evaluated using Wilcoxon test. j) ROC curve for OV RFU score as a predictive biomarker for stage-I HGOC vs benign patients.

We proceeded to test the performance of the 4 RFUs as a potential risk predictor for HGOC against benign ovarian tumors. We directly used the sum of downregulated RFUs (750 and 866) or upregulated RFUs (3808 and 1804) as predictors, and observed moderate predictive accuracy with an AUC slightly above 0.7. (Figure 3c-d). We combined the signals by using the up-subtracted by the down-regulated RFU sums, as “OV RFU score”. As expected, this score is significantly higher in the HGOC versus benign group (Figure 3e), with an improved AUC of 0.77 (Figure 3f).

To evaluate the reproducibility of this RFU score, we collected a validation cohort, which included 33 HGOC and 64 benign patients (Figure 3g, Table S3). All blood samples were collected before surgery for TCR-seq data generation. OV RFU score was able to separate HGOC from benign patients, but the predictive capacity was lower, with an AUC=0.66 (Figure 3h). However, unlike the discovery cohort, the validation cohort included 5 stage I HGOC patient samples (Table S3). We investigated the distributions of RFU scores within stage-I tumors and observed significantly higher scores than controls (Figure 3i). As a predictor, RFU score reached an AUC of 0.81 for stage-I HGOC vs control (Figure 3j). Interestingly, the scores of late stage HGOCs were lower than stage-I tumors, although statistical significance was not reached due to small sample size. These results indicated that the adaptive immune response may undergo nonlinear dynamic changes that peaks during the early progression of ovarian malignancies.

### Transient TCR repertoire changes in pre-diagnosis samples from ovarian cancer patients

The above findings hold promise in early ovarian cancer detection, yet further evaluation using more HGOC samples is challenging due to the rarity of stage I patients at diagnosis. To address this issue, we utilized blood samples collected from the Nurses’ Health Studies (NHS/NHSII). These studies, with over 280,000 participants, have collected blood samples from over 60,000 women primarily in the 1990’s and early 2000’s and followed women for diagnosis of ovarian cancer within 5 years after blood draw^29^. A subset of over 34,000 women gave two blood draws approximately 10-15 years apart. Among them, we identified 40 ovarian cancer patients (33 HGOCs) with two blood draws before diagnosis, one remote (≥10 years) and one recent (≤5 years). We also assayed 38 healthy controls matched on age at first and second blood draws (Figure 4a). All 156 NHS samples were sequenced for their TCR repertoires using the same commercial platform as the discovery and validation cohorts. We confirmed that within-individual dynamics is smaller than cross-individual variation^30^, with the 2^nd^ blood draw mostly similar to the 1^st^ draw from the same person (Figure S4). Given the higher RFU scores observed in stage I HGOCs (Figure 3i), we hypothesized that a transient change may occur in the adaptive immune repertoire within 5 years prior to the conventional diagnosis, when the tumor is still at an early stage.

**Figure 4.**
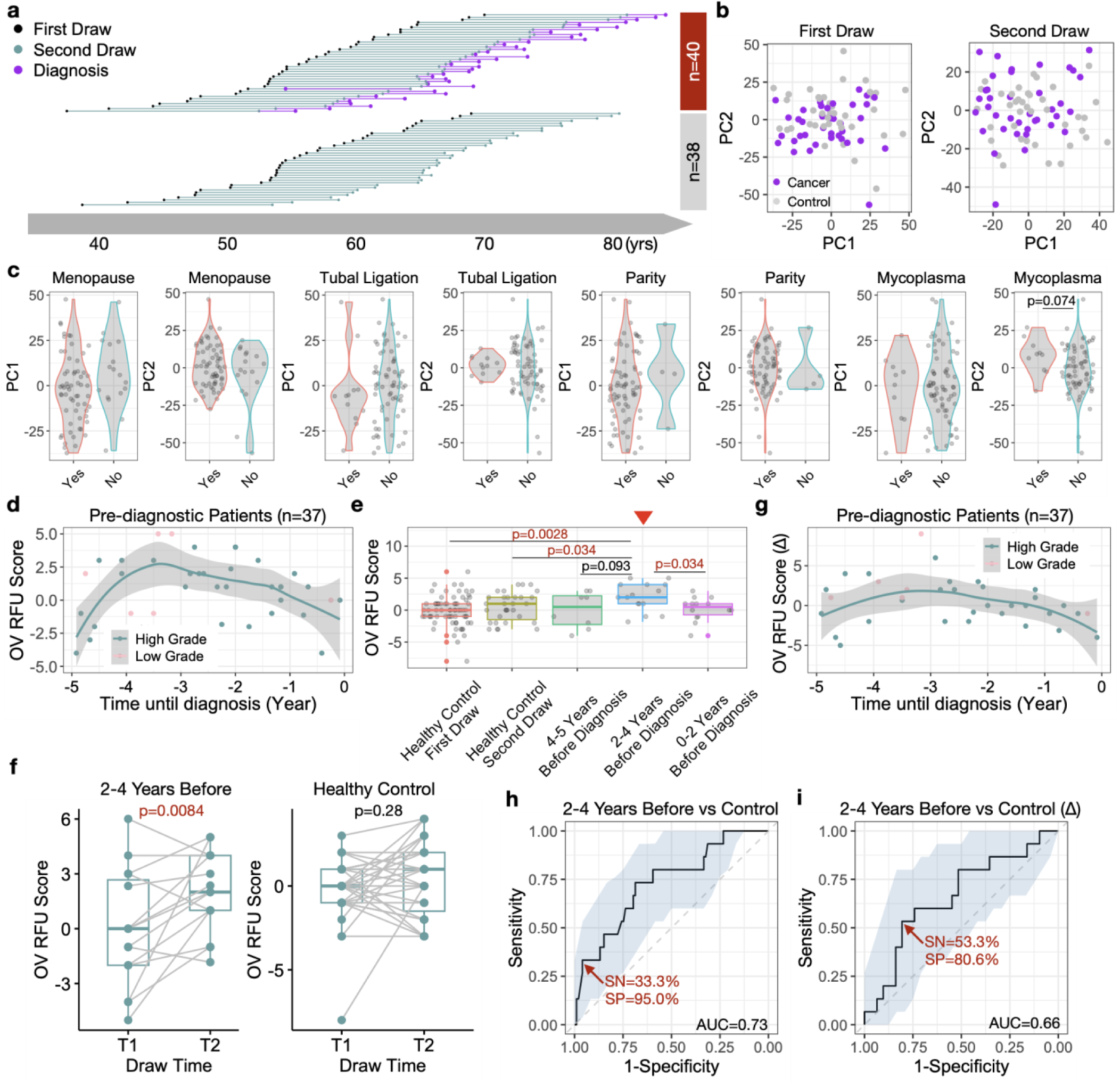
Dynamic changes of blood TCR repertoire prior to conventional ovarian cancer diagnosis. a) Diagram illustration of blood samples collected at first and second time point for both cancer and control subjects. b) PCA analysis of samples at first or second blood draws. c) Violin plot showing the distribution of PC1 or PC2 scores across putative ovarian risk factors. Statistical significance was evaluated using two-sided Wilcoxon test. d) Scatter plot showing the prediagnostic dynamics of OV RFU scores up to 5 years before conventional diagnosis. Loess smooth line was performed using only HGOC samples. Statistical significance was evaluated using permutation test. e) Boxplot showing the distributions of OV RFU scores in healthy controls and prediagnostic patients. Statistical significance was evaluated using two-sided Wilcoxon test. f) Paired boxplots showing the increments of OV RFU scores (2^nd^ timepoint – 1^st^ timepoint) in both patient and control samples. Statistical significance was evaluated using paired two-sample Wilcoxon test. g) Scatter plot showing the prediagnostic dynamics of incremental OV RFU scores. h-i) Prediction accuracy of OV RFU scores or increment scores for prediagnostic HGOC patients against healthy controls illustrated by ROC curves.

First, PCA plot of all samples at the first blood draw revealed no difference between cancer patients (10-15 years before diagnosis) and healthy control. At the second blood draw, there is a slight yet non-significant difference at PC2 (Figure 4b), suggesting that the cancer-induced changes were subtle, and may not drive the global alterations in the TCR repertoire. We next examined the impacts of known ovarian cancer risk factors^31,32^ on the immune repertoire. There was no difference between cancer patients and controls for menopausal status, tubal ligation, parity, or mycoplasma infection (Figure 4c). To avoid potential confounding effects, we removed all subjects with a family history of ovarian cancer in the downstream analysis.

We then proceeded to investigate the dynamics of OV RFU scores in the pre-diagnostic samples. RFU scores using the 4 RFUs described above were directly calculated for the 37 passed-filter ovarian cancer patients at the second timepoint. Interestingly, RFU scores displayed a significantly nonrandom dynamic curve prior to diagnosis (-5 to 0 years) that matched our expectation (Figure 4d). Specifically, the score rapidly increased -5 to -4 years, peaked around -3 years before slowly decreasing until the time of diagnosis. We dissected this period into three intervals based on the shape of the curve: uphill (-5 to -4), peak (-4 to -2) and downhill (-2 to 0). Direct comparison of the RFU scores of each group with the scores of healthy controls revealing that the peak group was significantly higher than both timepoints of the control cohort (Figure 4e).

Sampling two timepoints for both cancer and control cohorts allowed us to track the TCR repertoire changes over time. The RFU scores of patients within -4 to -2 years window showed significant increases when compared to their matched first timepoint, where the RFU scores of healthy individuals remained stable (Figure 4f). The increment of RFU scores between the two timepoints (Δ) displayed a nonlinear trend (Figure 4g). These results strongly supported our hypothesis that transient, but strong immune changes occurred during the early development of ovarian cancer. We therefore evaluated OV RFU score as a potential biomarker to detect ovarian cancer prior to its conventional diagnosis. If the disease is tracked within the -4 to -2 year window, the RFU score would reach an AUC of 0.73, with 33% sensitivity at 95% specificity (Figure 4h). The increment between timepoints (Δ) performed worse (Figure 4i). This is potentially because the random fluctuations in the TCR repertoire over 10 years reduced the signal/noise ratio.

## Discussion

In this work, we analyzed a total of 466 blood TCR-seq samples from ovarian cancer patients and healthy/benign controls. The computational analysis is based on a novel TCR embedding method specifically designed to quantify TCR repertoires. The RFU markers predicted in the discovery cohort were independently validated in two uniformly generated sample cohorts with age matched controls, thus avoiding potential batch effects or data leakage.

Mathematical models imply that TP53 mutations occurs several decades prior and precursor lesion STIC develops approximately 6 years prior to HGOC. Importantly, only a small fraction of individuals with TP53 mutations will develop HGOC and not all STIC progress to cancer. (Emerging evidence indicates that serous tubal intra-epithelial carcinoma (STIC), a putative precursor of HGOC, develops ∼7-12 years prior to diagnosis)^33^. Accordingly, the 1^st^ blood draw (>10 years) from the NHS cohort behaved similarly to the healthy controls (Figure 4b), while the 2^nd^ (<5 years) exhibited significant dynamic changes that strikingly coincides with the immunoediting process^17^. Specifically, during tumor initiation, the immune system functions well by priming T cells with tumor antigens, resulting in a rapid expansion of the tumor-reactive T cells in the lymphatic system, thus creating the ‘uphill’ part of the curve. Once the tumor cells reach equilibrium with the immune system, or the antigens have been sufficiently exposed to the T cells, a peak in the curve is anticipated. Finally, with tumor progression, immunosuppression becomes more dominant. T cell expansion slows down or stops with immune exhaustion^34^, potentially leading to a graduate decrease of tumor-reactive T cells in the blood. Despite these matched dynamics, it requires more clinical data to validate the pre-diagnostic behavior of the immune repertoire in ovarian cancer patients in larger sample cohorts and more diverse populations.

In this work, we selected four RFUs to construct the OV RFU score. It is unlikely that these RFUs encompassed all the TCRs informative to HGOCs. The number of markers discovered in this study was limited by the small number of HGOC samples, particularly early stage tumors. Further, all the HGOC patients in the discovery cohort were diagnosed at advanced stage, which, according to our observations above, may have yielded dampened immune response that might reduce statistical power. Although rare, a larger number of stage I HGOC samples with age-matched benign controls from future clinical studies will be ideal to uncover more informative RFUs and to improve the predictive accuracy. In addition, combining pre-diagnostic blood samples from multiple large prospective cohort studies could also increase power to identify novel early detection markers^35^.

We anticipate that 5,000 TCR clusters or RFUs do not completely cover the diversity of the immune repertoire. The TCRs captured and analyzed in this work are enriched for those with shared motifs across multiple individuals. Further, since the RFUs were defined using healthy donors, it is possible that disease-specific TCRs are underrepresented. TCR cluster analysis could be conducted on cancer patients to potentially identify RFUs that are specific to tumor antigens. Regarding ovarian cancer detection, with sufficiently more samples, RFUs can be redefined using HGOC patients, or even from ovarian tumor infiltrating T cells. HGOC-defined RFUs might further increase the prediction power when applied to prospectively collected patient cohorts.

Our study revealed interesting findings in the TCR repertoire of early-stage HGOCs, yet this work remains exploratory in nature. First, although the pre-diagnostic curve of RFU scores matched our findings in the validation cohort of cases and benign controls, there is no definitive evidence to support that women with samples collected at the ‘peak’ phase had *bona fide* stage I HGOCs. Next, our analysis was performed at the RFU level, where the antigen-specificity of individual TCRs was not investigated. This is mainly due to the lack of established T cell antigens from early-stage HGOCs, which can be improved with future immunogenomic research on ovarian cancers. Finally, as a screening biomarker, the sensitivity and specificity of the RFU score are far lower than what would be required to reach 10% positive predictive value^8^. As a diagnostic tool, it is less accurate than the established indices, such as ROMA^36^ or RMI^37^. As discussed above, this limitation might be addressed by including more TCR-seq samples. Due to the rarity of stage I HGOCs, future emphasis could be given to large clinical networks to recruit such patients or that have biobanked such samples^38^.

Despite these limitations, our analyses support that ovarian tumor progression causes observable changes in the blood TCR repertoire, which are stronger at early stage when HGOC is immunogenic and more likely to be curable. It further suggests that studies in higher stage tumors, where immunosuppression is more common, may not be warranted. In addition, our study demonstrates the value of using prospectively collected samples in identifying new biomarkers through quantification of these changes in RFUs, which may lead to practical solutions for immune-based ovarian cancer early detection.

## Author contributions

B.L. and J.L. conceived the project. B.L. developed the RFU method and performed computational analysis with X.Y. J.L. lead sample collection at UTSW. C.A.H. and S.T. provided NHS samples and associated clinical data. J.Y. performed gDNA extraction and sample QC. B.L. wrote the manuscript with J.L. B.L. and J.L. supervised the study.

## Acknowledgement

This work is supported by NCI R01 grants CA258524 (B.L. and J.L.) and CA245318 (B.L.). Sample collection efforts received support from the Department of Obstetrics and Gynecology, UT Southwestern Medical Center.

## Data and code availability

R codes for RFU calculation are available at Github: https://github.com/s175573/RFU. Due to concerns regarding patient privacy and data sharing policy of Nurses’ Health Study, the raw data will be available upon request under appropriate IRB protocol. In the meanwhile, the processed RFU matrices necessary for repeating the results of this work are available at: https://github.com/s175573/RFU/tree/main/Data

## Methods

### Description of TCR repertoire samples and preprocessing

All TCR repertoire sequencing samples that were not produced from this study were accessed from the immuneAccess database managed by Adaptive Biotechnology. These samples were profiled using the immunoSEQ platform developed by the company. Zip files were directly downloaded through the ‘Export’ function and selecting ‘v2’. Accession numbers for each cohort are available in Table S1. For each repertoire sample, sequences with missing variable genes or nonproductive CDR3 regions were removed. The top 10,000 TCRs with most abundant clonality were selected for RFU calculation. These preprocessing criteria were applied to all the TCR-seq samples throughout this study.

### Ovarian cancer patient cohorts and Nurses’ Health Study samples

Women present with ovarian tumors were consented at Parkland Hospital at UT Southwestern Medical Center (UTSW) between 2019-2022. No stage-I HGOC patients were collected during this period. Blood samples were collected prior to surgeries and stored in EDTA tubes in -80 degree freezer. Tumor histology, including benign or malignant, was available after pathological verification. Sample collection was approved by the Institutional Review Boards (IRB) with protocol number STU-2020-442. All samples collected at UTSW were used as the discovery cohort. Buffy coat samples of patients in the validation cohort were purchased from Accio Biobank Online in 2021.

Additional samples were obtained from the Nurses’ Health Studies (NHS/NHSII), two large prospective cohorts starting in 1976 (NHS) and 1989 (NHSII), with over 238,000 women. Between 1989 and 1990, 32,826 NHS participants donated self-collected blood samples, which were shipped on ice via courier where and processed into plasma, red blood cell, and white blood cell components; a second collection in 2000-2002 from over 19,000 of these women used similar protocols. Similarly, between 1996 and 1999, 29,611 NHSII participants donated blood samples; a second collection occurred from 2011-2014. Cases of ovarian cancer were identified via self-report on biennial questionnaires, report of family members, or via the National Death Index. Medical records or reports from cancer registries were used to confirm the diagnosis. Cases were matched to controls were were alive and had at least one ovary at the time of the case diagnosis and matched on age, menopausal status, date and time of blood collection, fasting status, and hormone therapy use. For this analysis, we assayed both the first and second blood draw samples from cases that were diagnosed within 5 years after the second blood draw and their matched controls. De-identified patient information, including age at blood draws, age at diagnosis, tubal ligation status, parity, menopausal status, and other ovarian cancer risk factors were provided for the analysis.

### Genomic DNA isolation and TCR repertoire sequencing

Genomic DNA was isolated from 200 μl whole blood (from UTSW) or 10 ul buffy coat (from Accio Biobank or NHS) using the DNeasy Blood and Tissue Kit (Cat# 69504, Qiagen) following the manufacturer’s guidelines. gDNA concentration was measured using a NanoDrop 2000 Spectrophotometer (Thermo Fisher Scientific). The purity of gDNA was determined by measuring the 260-280 nm absorbance ratio. Optimal purity was expected to be in the range of 1.7-2.0. The integrity of the gDNA samples was assessed for evidence of degradation using agarose gel electrophoresis. Appropriate quality gDNA was expected to migrate predominantly above 10 kb on agarose gels. All samples passed DNA purity and integrity quality controls. Twenty samples of gDNA were sent to Adaptive Biotechnology for targeted TCR β chain repertoire sequencing using immunoSEQ at survey sequencing depth. Raw TCR reads were processed with immunoSEQ Analyzer for CDR3 assembly, variable/joining gene calling, and clonal frequency estimations.

### Repertoire Functional Unit method description

#### i) TCR embedding

We applied GIANA^19^ to perform clustering of over 20 million TCRs using both the CDR3 sequences and variable gene alleles obtained from public domain (Figure 1b). These samples covered a wide spectrum of disease context, including healthy individuals and patients with cancer, autoimmune disorders as well as viral infections (Table S1). Previous work, including ours, have demonstrated that TCRs clustered using such strategy are highly specific (≥95%) to the same antigen epitopes^21–23^, with smaller (n≤5) clusters being more likely to share antigen-specificity^19^.

From GIANA output, we identified a total of 821K such clusters. An example of a typical cluster of two sequences, CSARQG*<u>A</u>***<u>R</u>***<u>T</u>*YEQYF and CSARQG*<u>A</u>***<u>Y</u>***<u>T</u>*YEQYF, bear a mismatch R/Y in position 8 (Figure 1c). We considered the amino acids flanking this mismatch and extracted the trimer sequences from both TCRs. As the two TCRs likely share antigen-specificity, the two trimers, ART and AYT, are thus considered ‘replaceable’ in the context of antigen recognition. We then traversed all 821K clusters and built the 8,000-by-8,000 trimer-substitution matrix (TSM) by calculating the number of replacements of each trimer pairs (Figure 1d). We calculated the Spearman’s correlation matrix using TSM and converted it into a Euclidean distance matrix (EDM). Next, similar as in GIANA, we obtained the isometric embedding vector for each of the trimers using multi-dimensional scaling based on the EDM. This approach allowed us to use a numeric vector to represent each trimer, with similar trimers located closely in the Euclidean space (Figure 1e). The embedding of each CDR3 sequence is then calculated as the average of all the vectors from consecutive trimers (Figure 1f). This embedding is a continuous representation of TCR similarity.

#### ii) Benchmark using antigen-specific TCRs

We benchmarked the trimer-based embedding using 1,031 TCR sequences with known antigen-specificity to 9 epitopes (Table S2). This dataset has been used in our previous work to benchmark the specificity of TCR clustering^23^. To avoid bias towards epitope(s) with excessive amount of TCRs, we restricted the antigens with <170 TCRs. Coordinates were calculated for each CDR3 sequence. For each pair of TCRs, we calculated the Euclidean distance as the predictor, with the response being if or not the two TCRs share the same antigen. From the total of half million comparisons, we excluded pairs with distances >0.025, based on the fact that trimer-based embedding is only powerful to identify shared antigen specificity among TCRs with similar sequences. ROC curve was generated with the above predictor and response variables, with AUC calculated using the curve.

#### iii) Definition of Repertoire Functional Units (RFU)

We pooled 1.2 million TCRs from 120 healthy donors from a previous study^25^, and projected them onto the Euclidean space with trimer-based embedding. We divided the TCR sequences in this space into 5,000 groups with the k-means method. We referred the centroid of each group as a ‘Repertoire Functional Unit’, or RFU. To calculate the RFU vector of a new TCR repertoire sample, we first select the top 10,000 most abundant TCRs based on clonal frequencies. For each TCR, we calculate the embedding vector and assign it to the closest centroid from 5,000 RFUs. The value of each RFU is determined by the number of TCRs assigned to its centroid. We chose 5,000 as the group number so that the expected count for each RFU is 2.

### Statistical Analysis

Computational and statistical analyses in this work were performed using the R programming language v4.3.0. Logistic regression adjusted for patient age and race (Figure 2) was implemented using the *glm* function. FDR control was using the Benjamini-Hochberg method. Sequence logos were generated using package *ggseqlogo* (v0.1), by performing multiple sequence alignment (*msa*, v1.32.0) using CDR3s with length 16. Donut plot (Figure 3) was generated using package *webr* (v0.1.5). ROC curves with 95 confidence intervals and AUC values were generated using package *pROC* (v1.18.2). Neighbor joining trees were calculated and visualized using R package *ape* (v5.7-1). Subpanels of main figures were produced using *ggplot2* (v3.4.2). Permutation test in Figure 4d was performed as follows: with the goal of testing how significant the peak-like dynamics of prediagnostic curve, we randomly permuted the RFU scores for 1,000 times and recalculated the Loess smooth curve with default parameters (R function *loess*). For each permutation, we calculated the range of the curve (max - min), denoted as D_r_. The range of the unshuffled curve is denoted as D_0_. p value was estimated as the number of permutations with D_r_ greater than D_0_ divided by 1,000.

## Supplementary Figure Legends

**Figure S1.**
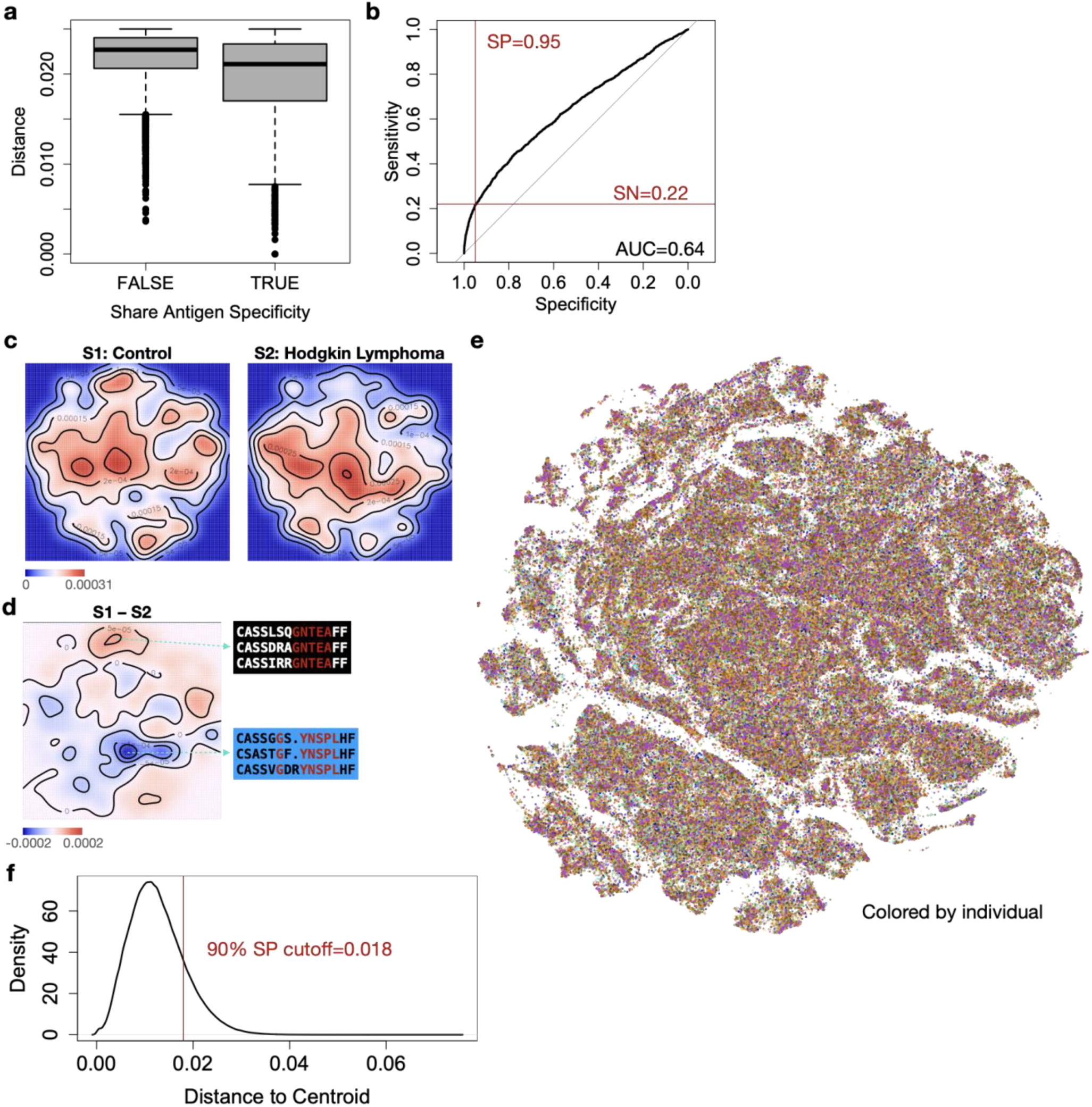
Benchmark of trimer-based embedding and repertoire functional units. a) Boxplot showing the distributions of Euclidean distances between a pair of TCRs with known antigen specificities in the benchmark dataset. b) Prediction accuracy of Euclidean distance on if or not the pair of TCRs sharing specificity by ROC curve. c) 2-D density plot showing TCR distributions in a healthy control and a Hodgkin lymphoma patient. d) Density difference from c) showing the enriched or depleted regions in the TCR embedding space. Selected TCR motifs were associated with these regions. e) Same t-SNE plot as in Figure 1g, except colored by different individuals. f) Distribution of Euclidean distance between any TCR to its assigned k-means cluster centroid. 90% specificity cutoff was determined by the ROC curve in b).

**Figure S2.**
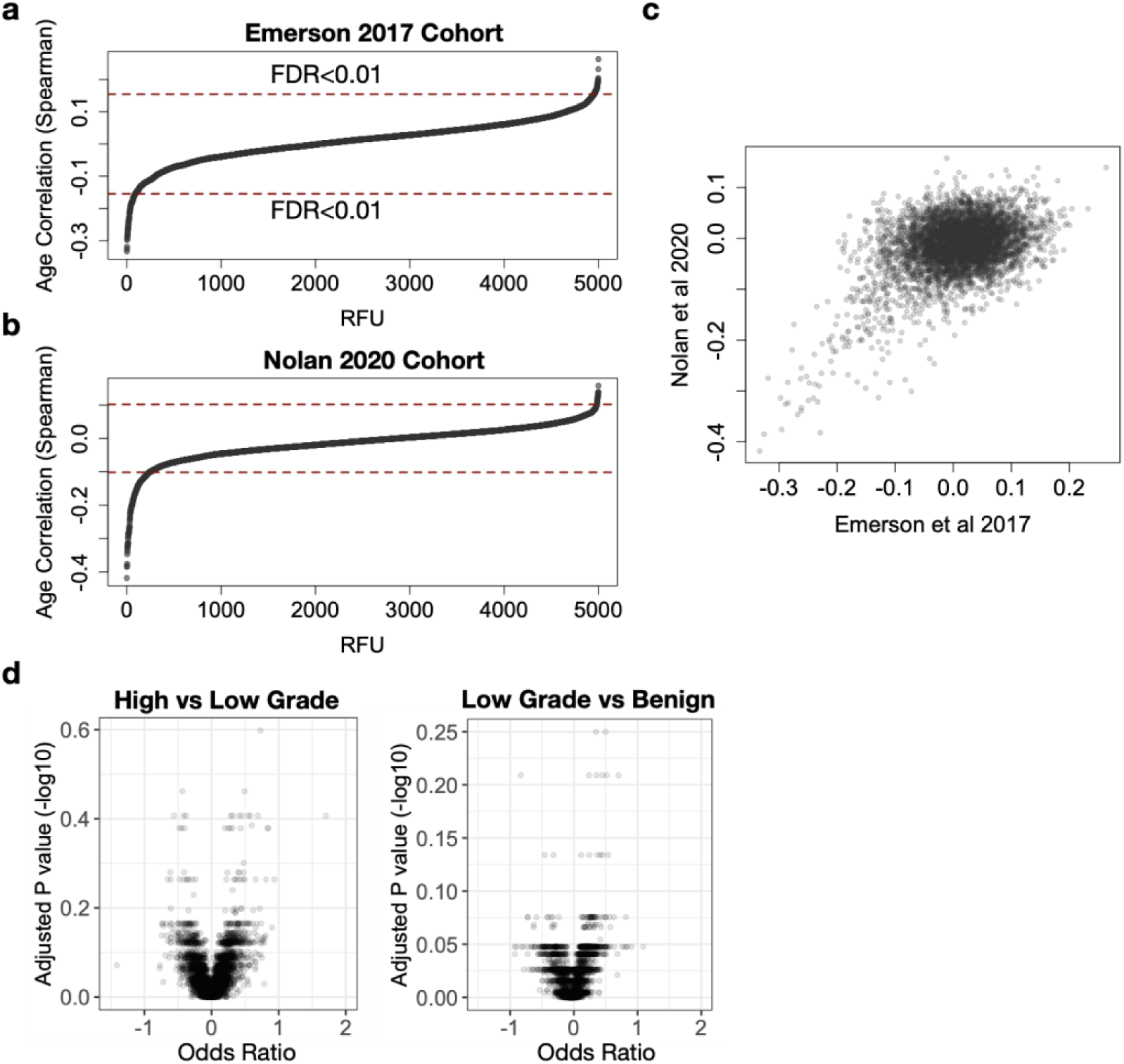
Additional analysis related to ovarian cancer samples collected in the discovery cohort. a-b) Ordered Spearman’s correlations of age and each of the 5,000 RFUs with dashed red lines marking FDR<0.01. c) Scatter plot showing the relationships of age associations for each RFU between the two large healthy donor cohorts in a-b). d) Volcano plots showing the output of the same analysis as in Figure 2f for the other disease categories in the discovery cohort.

**Figure S3.**
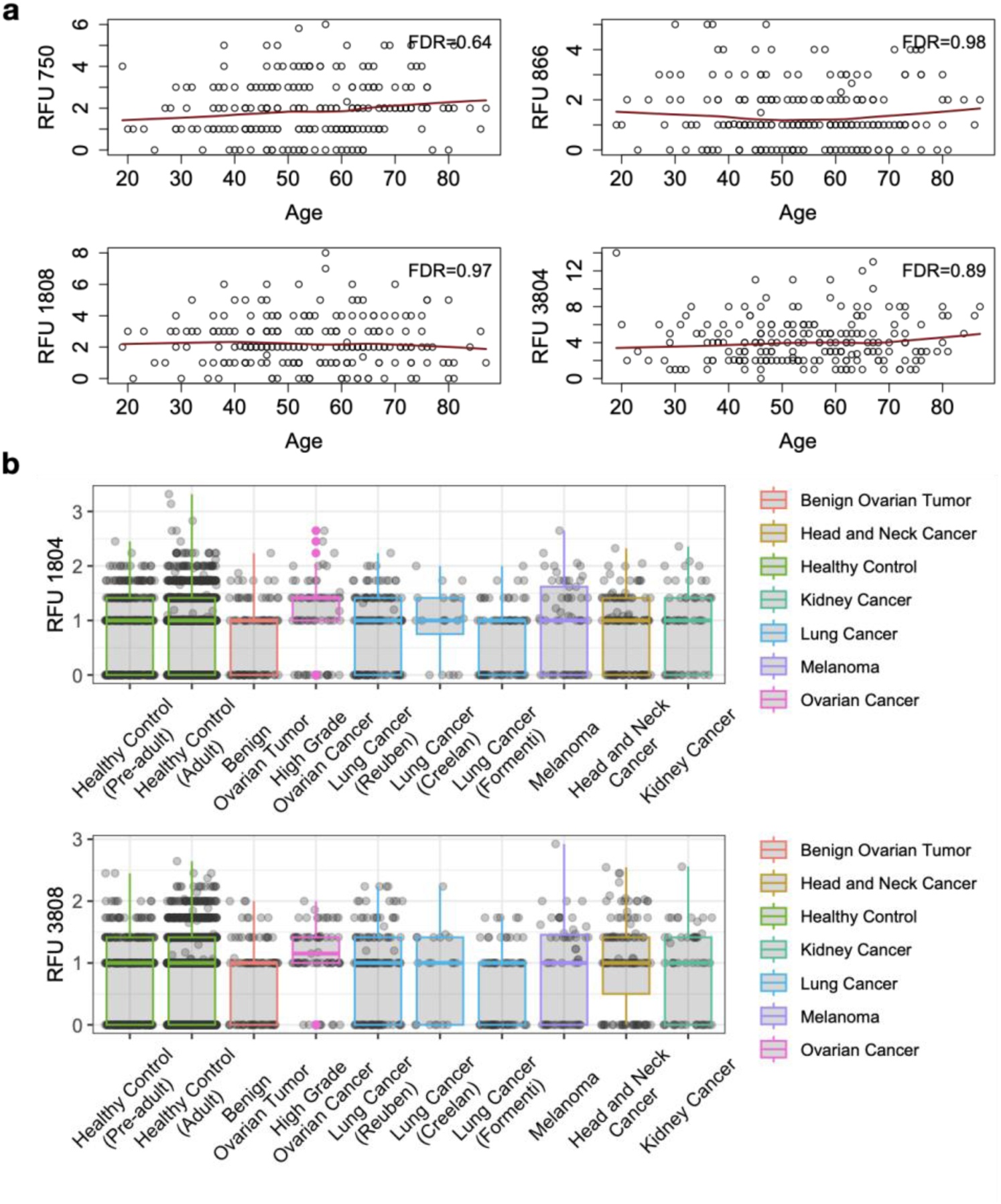
Distribution of selected RFUs across age or in the TCR repertoires of multiple cancers. a) Age association of selected RFUs in Figure 3a. Loess smooth curve was shown as red line in each panel. Spearman’s correlation test was used to evaluate statistical significance and FDR was adjusted using the Bejamini-Hochberg method across all 5,000 RFUs. b) Same analysis as in Figure 3b showing the distributions of the upregulated RFUs in the top list.

**Figure S4.**
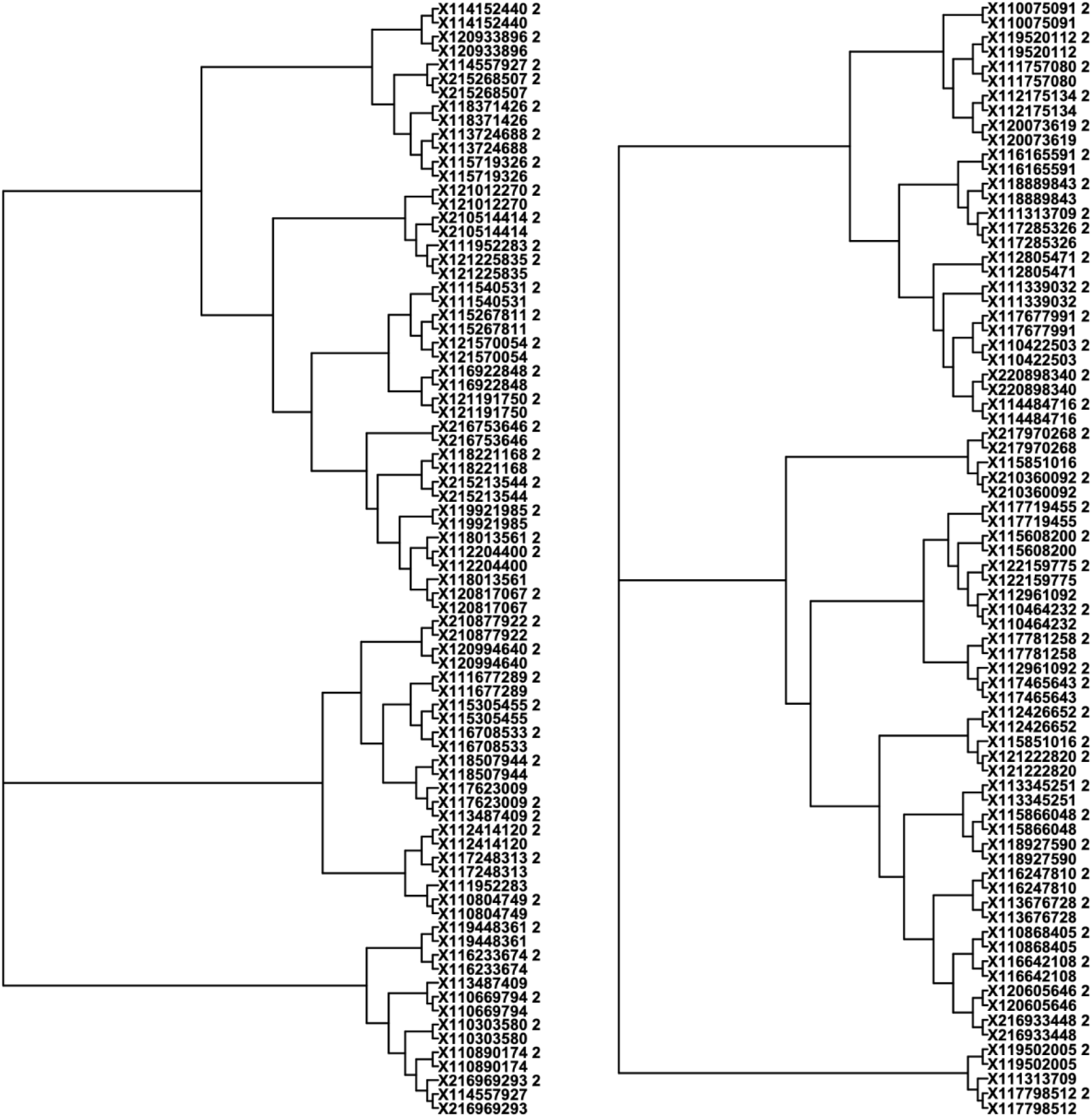
Neighbor-joining tree of TCR repertoire samples from NHS cohort. Distance matrix was calculated as squared root of 1-Spearman’s correlation for the patient (left) and donor (right) samples separately. Neighbor joining trees were generated using the distance matrices. The second timepoints were marked with ‘2’ at the end of the label.

## Supplementary Table Legends

Table S1. TCR repertoire sample cohorts used to generate the clusters for trimer substitution matrix.

Table S2. Benchmark data set of TCR sequences with known antigen epitopes.

Table S3. Summary of ovarian cancer samples collected in the discovery and validation cohorts.

Table S4. Demographic information of samples in the NHS cohort with RFU scores included.

